# Iterative feature selection method to discover predictive variables and interactions for high-dimensional transplant genomic data

**DOI:** 10.1101/605428

**Authors:** Hu Huang, Cynthia Vierra-Green, Stephen Spellman, Caleb Kennedy

**Author notes:** Corresponding author: Hu Huang.

## Abstract

After allogeneic hematopoietic stem cell transplantation (allo-HCT), donor-derived immune cells can trigger devastating graft-versus-host disease (GVHD). The clinical effects of GVHD are well established; however, genetic mechanisms that contribute to the condition remain unclear. Candidate gene studies and genome-wide association studies have shown promising results, but they are limited to a few functionally derived genes and those with strong main effects. Transplant-related genomic studies examine two individuals simultaneously as a single case, which adds additional analytical challenges. In this study, we propose a hybrid feature selection algorithm, iterative Relief-based algorithm followed by a random forest (iRBA-RF), to reduce the SNPs from the original donor-recipient paired genotype data and select the most predictive SNP sets in association with the phenotypic outcome in question. The proposed method does not assume any main effect of the SNPs; instead, it takes into account the SNP interactions. We applied the iRBA-RF to a cohort (*n*=331) of acute myeloid leukemia (AML) patients and their fully 10 of 10 (HLA-A, -B, -C, -DRB1, and -DQB1) HLA-matched healthy unrelated donors and assessed two case-control scenarios: AML patients vs healthy donor as case vs control and acute GVHD group vs non-GVHD group as case vs control, respectively. The results show that iRBA-RF can efficiently reduce the size of SNPs set down to less than 0.05%. Moreover, the literature review showed that the selected SNPs appear functionally involved in the pathologic pathways of the phenotypic diseases in question, which may potentially explain the underlying mechanisms. This proposed method can effectively and efficiently analyze ultra-high dimensional genomic data and could help provide new insights into the development of transplant-related complications from a genomic perspective.

## Introduction

Acute graft-versus-host disease (acute GVHD) is one of the major complications after HLA-matching allogeneic hematopoietic stem cell transplantation (allo-HCT) that cause non-relapse morbidity and mortality, affecting up to 40∼60% of transplant patients and accounting for 20% of deaths after allogeneic HCT. It is an immunologically mediated complex disease. To date, genome-wide association studies (GWAS) and candidate gene studies have identified SNPs associated with acute GVHD, including SNPs that cause the genetic disparities between the donor and the patient, i.e., the minor histocompatibility antigen (MiHA) single nucleotide polymorphisms (SNPs)[1], and SNPs that modify gene functions [2]. However, the genetic risks for acute GVHD outcome have not been well defined yet [3]. Most such studies have focused on single locus variants individually or a few candidate gene locations and tested them for association with acute GVHD. Unlike the assumptions of these studies, however, genes tend to interact within specific regulatory and functional pathways, contributing to the disease development.

Next-generation sequencing technologies have enabled affordable high-throughput whole genome microarray genotyping and sequencing. These technologies pose multiple unique challenges in transplant-related genomic studies that need to be addressed and taken into consideration. First, each allo-HCT case involves in two individuals, the donor and the patient, both of whose genomes directly influence the transplant outcomes. Thus, the genomic association models should consider two genomes simultaneously as a single ‘sample,’ whereas, in common disease association studies, either the donor or recipient genome is considered as a single sample. Second, the transplant-related outcomes are caused by the genomic disparities between donor and recipient with their synergistic interactions, and hence there is no inheritability of the diseases. Third, the allele frequencies may not play an as much of an important role as in the common disease association studies; instead, the combinations and mismatches of donor-recipient (DR) pair genotypes may be more influential. Fourth, the cohorts in the transplant genomic studies are more heterogeneous and harder to control than in common disease studies. Each year, there are limited transplant cases due to the challenges of finding HLA-matching unrelated donors and hence it is harder to recruit groups that share most of the conditions. Furthermore, the cohort size usually is very small compared to the common disease studies, and this also leads to the lack of publicly available transplant-related genomic databases.

Alloimmune complications after transplantation, such as acute GVHD, not only involve immune responses to conventional exogenous antigens but also responses to alloantigens. The latter is unique to transplant cases. The major player in GVHD is the activated T cells that recognize and eliminate alloantigens. These T cell functions are influenced by the complex interactions between regulatory networks, pathways, extracellular environment and the unique conditions induced by transplantation procedures [4]. Thus, it is reasonable to assume that both donor’s and recipient’s genomes matter in the development of acute GVHD. However, most transplant-related outcome studies often focus on patients’ genomes, and very few studies have examined both HLA-matching donor and recipient genomes together [5]. Here, we assume the donor’s genome as equal weight as the recipients and form a paired genotype encoding matrix from each transplant case. With a sufficiently large sample size and appropriate models, we can capture the interacting signals from the paired genome.

Similar to the general whole-genome research in common disease studies as Moore and Ritchie outlined [6], transplant-related genomic research also faces three major challenges. The first challenge is to identify meaningful genetic variants along with clinical characteristics that are susceptible to transplant-related complications. The genetic variants include SNPs, genes, or specific gene regions. As described above, transplant-related complications are mostly caused by the genetic disparities between donor and recipient and the combination of their clinical and demographic characteristics (e.g. sex, age, race, and ethnicity), rather than the disease heritability. The second challenge is to build robust and powerful predictive models that take both genetic and demographic variables into account and output the probability of developing adverse transplant outcomes given a candidate graft characteristic. The predictive models will help facilitate effective and optimized donor search strategies with the best transplant outcomes. While the first two challenges are from statistical and machine learning aspects, the last challenge is to interpret the genetic variants and the predictive models from a biological perspective and further advance our understanding of the transplant-related complications. Biological functional interpretation will help optimize the donor selection process, improve the transplant outcomes and prevent transplant-related complications. It is the most important and difficult challenge and requires a deep understanding of human immunology as well as genetic regulatory mechanisms. Wet lab bench experiments would be the most effective way to validate the hypotheses but it would be too time-consuming and could become impossible if there are too many factors to control. It is one of the current leading translational bioinformatics research focus areas.

Traditional logistic regression models, *χ*^2^-test, and odds-ratio are efficient and intuitive when finding simple linear relationships from a large-scale data set; however, they have limited power in modeling high-order non-linear relationships among variables, especially for ultra-high dimensional data. Whole genome microarray genotype data usually cover over 500,000 base pairs of genetic variables and a majority of them may be considered as noise since they do not show any susceptibility to the diseases in question. Data mining or machine learning techniques build models without any linearity assumptions on the data and can identify the high-order interactive relationship among variables. This is especially attractive to genomic data mining tasks. From a machine learning point of view, there are two main tasks in this context: 1) select the most informative variables from the over 1 million SNPs; 2) predict the disease risk from the selected variables using classifiers. From a clinical point of view, these selected variables should be interpretable. Unlike Mendelian diseases, transplant-related outcomes are influenced by non-linear interactions of multiple genes between donor and recipient. Transplant-related outcomes are more likely a joint effect of multi-factors rather than one single main effect factor. The attribute or feature interaction methods in machine learning seem more appropriate in this case. The data-mining methods can detect nonlinear relationships that traditional regression-based models cannot represent, and this is especially true for dealing with high-dimensional data. In addition, the data-mining algorithms may also uncover the interactions between variables other than their main effects. Applications of machine learning in detecting gene-gene interactions in genetic epidemiology are reviewed in [7–9].

The purpose of this study is to investigate the application of machine learning techniques in transplant genomics. More specifically, we propose a hybrid feature selection model (iRBA-RF) by incorporating the iterative Relief-based algorithms (iRBA) and a random Forest (RF).

The rest of the paper is organized as follows. First, we define the transplant genomics and outcome association study in the machine learning context. Second, we briefly review feature selection and classification models. Then we apply the proposed iRBA-RF model to transplant cases to identify critical genetic factors. Lastly, we show the predictive results and provide a possible biological interpretation, as well as the applicability, limitations and future work.

## Methods

### Problem Definition

In allo-HCT, histocompatibility of stem cells is the primary concern of graft selection, and there are many factors involved in the donor screening process. In this study, we retrospectively investigated HLA-A, -B, -C, -DRB1, and -DQB1 fully matched (10/10) unrelated donor transplant cases, and explored the potential genetic variants that may influence the transplant outcomes. In addition to minor histocompatibility antigens (MiHAs), there are other genes involving in regulatory immunological pathways that are critical to the development of GVHD. In complex diseases, there is overwhelming evidence that non-additive synergistic effects of multiple genetic factors play an essential role in the development of the diseases. As described before, we consider the donor genome the same weight as the recipients.

In order to investigate the applicability of the proposed model in the transplant-related genomic studies, we assess the following two case-control scenarios: 1) Scenario 1 (AML case-control): acute myeloid leukemia (AML) patients as case and their HLA-matched healthy donors as the healthy control; 2) Scenario 2 (aGVHD case-control): the donor-recipient (DR) pairs where the patients developed the acute GVHD symptoms as the case and the DR pairs where the patients did not show any adverse symptoms as the controls.

The main difference between these two scenarios in the context of machine learning is how the genotypes are represented as a feature matrix. Scenario 1 is a common case-control situation where each individual’s genotype vector is a single observation, and the AML disease condition is the phenotypic outcome to be predicted. In Scenario 2, an observation is defined as the combined genotype vectors of the recipient and the donor, where the length of the vector is doubled compared to Scenario 1. In addition to the DR genotypes, other clinical characteristics may be included in the model, such as the HLA typing and the donor-recipient sex-mismatch status.

### iRBA-RF: a hybrid feature selection model for detecting attribute interactions

In bioinformatics, the “large *p* small *n*” problem is a common challenge, especially when it comes to genomic association analysis. The most common problems in genomics data are 1) noisy data 2) heterogeneous data types and 3) ultra-high dimensional feature space. In machine learning, the feature selection procedure is employed to avoid the “curse of dimensionality” for small samples with high dimensions [10–12]. The objective of feature selection is to select the most relevant feature subset to achieve the best classification/prediction performance without losing the generalization power (accuracy, speed, and generalization). A strong feature relevance indicates the feature is necessary for the predictive model, while an irrelevant feature does not contribute to the predictability. In some cases, the presence of certain features would decrease the predictability of the model, in which case they are considered as noise. For a formal theoretic derivation of feature relevance, interested readers may refer to [13].

Depending on the feature search strategy and the level of predictive classifier integration, there are three different categories of feature selection methods: filter, wrapper and embedded. Filter approaches are independent of classifiers; instead, they examine the intrinsic properties and relationship between the phenotype in question. Specifically, the information theoretic metrics, such as mutual information [14, 15] and entropy/information gain [16, 17], are popular options to measure the intrinsic properties. Since these approaches do not involve training a classifier, they are computationally fast and applicable to a large dataset. Detailed reviews of feature selection techniques in bioinformatics can be found in [18, 19].

Since we are interested in interpretable variables that are linked to the phenotypes within a reasonable computation time, we adopt the filter-based approaches. More specifically, we propose a hybrid feature selection model that combines an iterative Relief-based algorithm and a random forest (iRBA-RF), to iteratively eliminate the irrelevant features and select the top-ranked features, respectively. In the next subsections, we describe the details of each algorithm.

### Iterative RBA for variable elimination

The Relief-based algorithms (RBAs) was inspired by instance-based learning [20, 21], where it draws instances at random and iteratively compute and updates the weights of features based on their nearest neighbors and their phenotypes. The features that distinguish the selected instance from its neighbors of a different class get more weight. The original Relief algorithm only compares one nearest neighborhood of each class, which is sensitive to noisy data and restricted to a binary classification problem. There have been many studies to address the limitations and improve the performance of the original Relief algorithm. The most widely used RBA is ReliefF [22], which relies on the nearest *k* neighborhoods, instead of one. By comparing the entire vector of values across all attributes among neighbors, ReliefF can capture the attribute (feature) interactions and has gained popularity in data mining applications. Figure 1 shows an example of ReliefF on acute GVHD outcome data set with *k* =3 nearest neighbors in each class, respectively.

**Figure 1.**
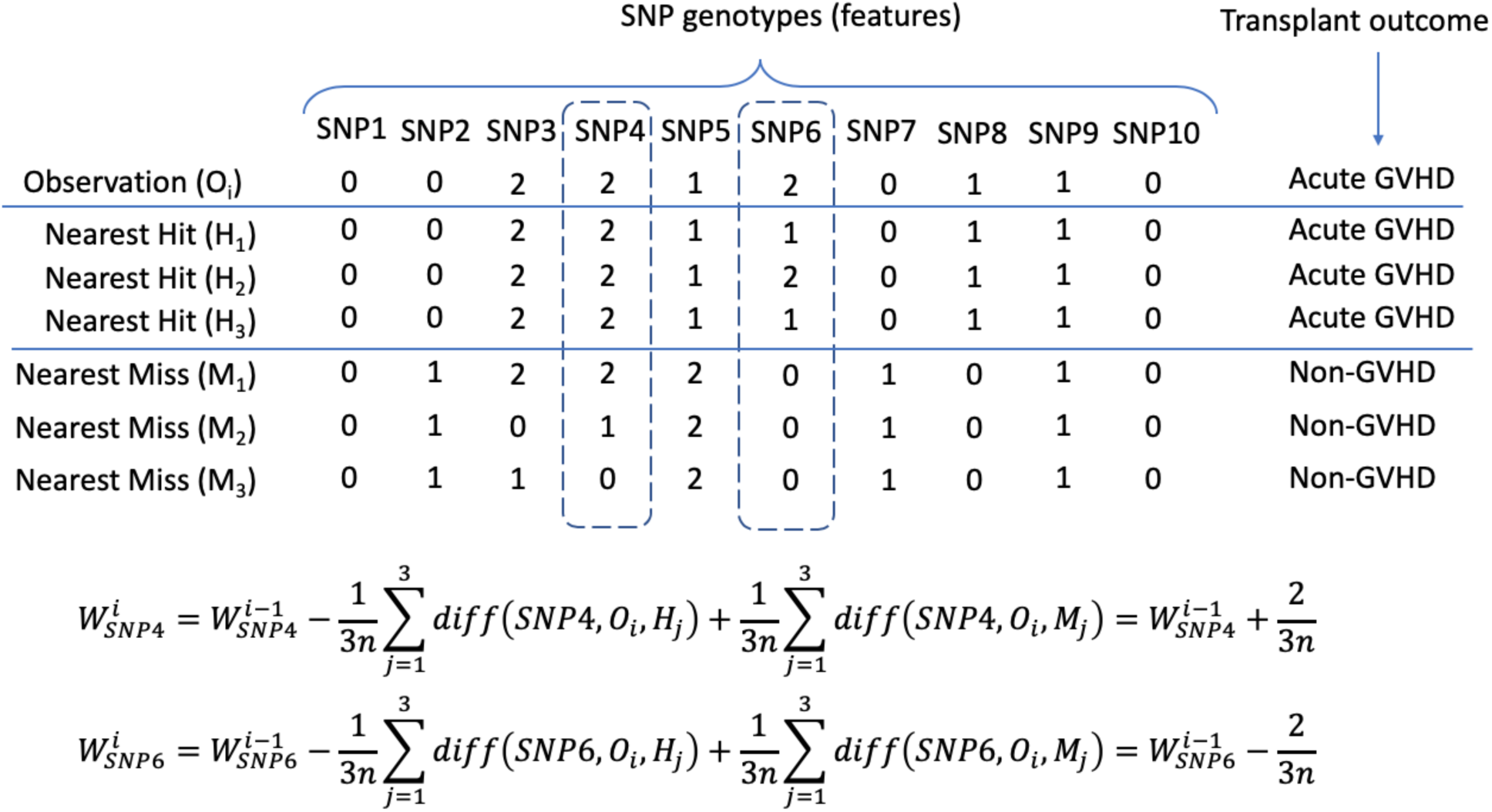
Illustration of ReliefF algorithm with *k*=3 nearest hits and misses, respectively, on transplant outcome data.

However, ReliefF is not robust to noisy features where it cannot capture the correct signal. An improved ReliefF called Tuned ReliefF (TuRF)[23] was proposed to iteratively remove features that have low-quality scores, in most cases are noisy features. More extended RBAs were later developed and applied in genomic data analysis, including Spatially Uniform ReliefF (SURF)[24], SURF*[25], SWRF*[26], Multiple-Threshold* (MultiSURF*)[27], and MultiSURF[28]. They use different strategies to select neighboring hits and misses and calculate their weights to improve sensitivities and computational efficiency. Furthermore, unlike the original Relief algorithm, these improved versions can handle incomplete data and extend to multi-class problems. For an in-depth review of RBA-based feature selection methods, readers may refer to [29].

In typical genomic association studies, there are over 500,000 SNPs to be examined. Especially in the context of transplantation, donor-recipient pair genotypes may include over 1 million SNPs. This poses a challenge in computational efficiency. For such ultra-high dimensional genomic data, iterative and efficient approaches that are wrapped around and integrated into the above core RBAs are recommended. VLSReliefF [30] algorithm is reported to be able to detect feature interactions in very large feature space both efficiently and accurately. The main idea is to randomly group ***s*** subsets of the feature set with ***a***_***s***_ features and individually apply ReliefF to each group to calculate local feature weights. The global weights of each feature are the maximum value of the local feature weights among the subsets. In this study, we follow the framework of VSLReliefF and repeat the process multiple times to remove low-quality features iteratively, as shown in Figure 2. Instead of ReliefF, here we choose MultiSURF as the core RBA since it has shown to outperform in multi-way interaction detection as well as various associations, compared to the other RBA algorithms [28].

**Figure 2.**
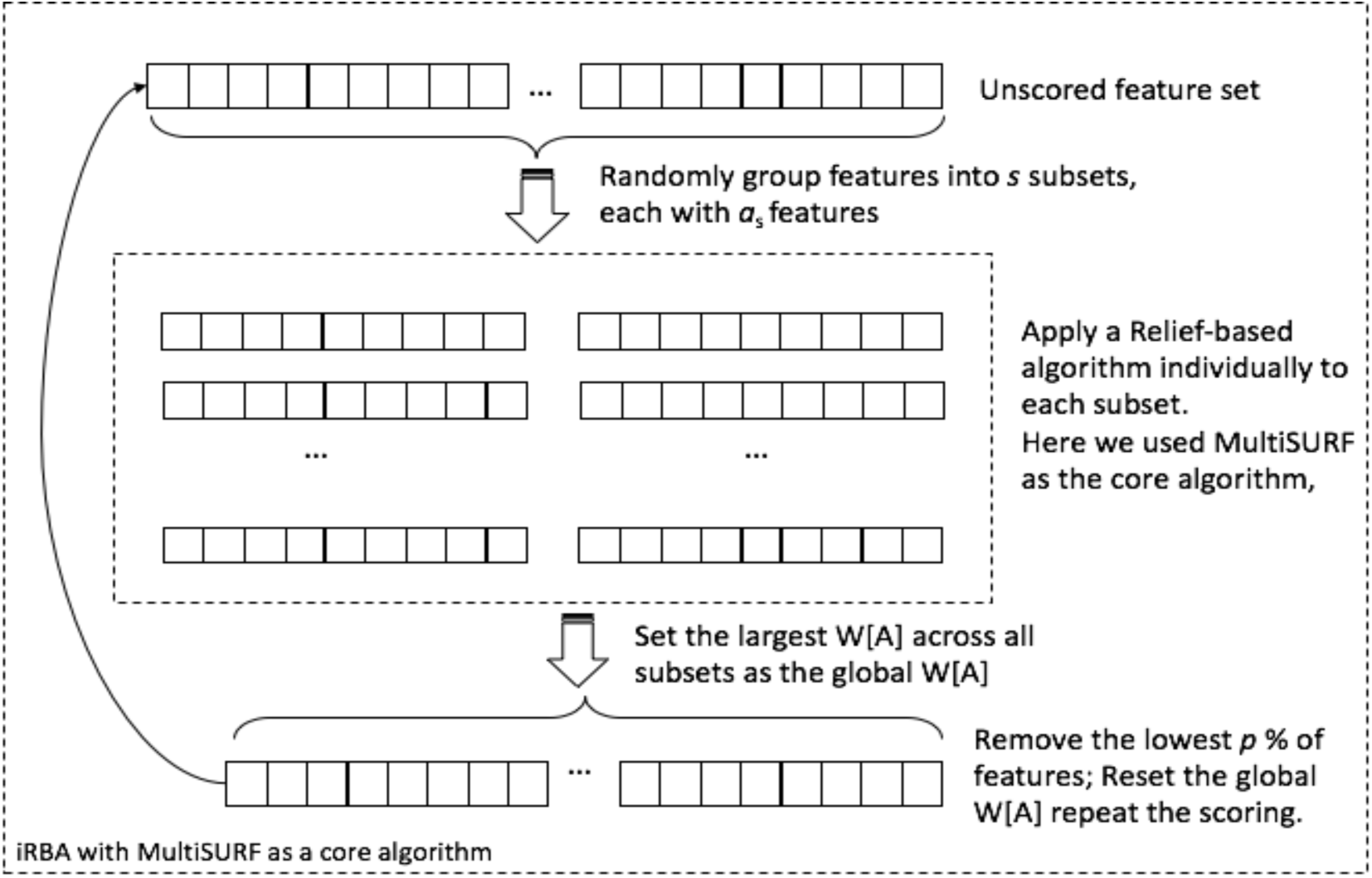
Illustration of iRBA, adapted from [29].

### Random forests for feature importance ranking and variance selection

Random forests (RF) are ensembles of tree-structured classifiers that are constructed in the following random fashion: each tree is grown using a bootstrap sample, i.e., aggregated sampling with replacement, of original training set and a randomly chosen subset of features and a majority voting scheme to ensemble individual trees, as illustrated in Figure 3 [31]. Instead of using the whole set of a training set, each tree is trained on the bootstrapped sample set, and the rest samples are used as a validation set to estimate the tree’s classification error. This validation set is called the out-of-bag (OOB) samples. The OOB scheme is used to monitor the generalization error, strength, and correlation of trees in the forests, as well as the variable importance. As more trees added to the RF, it is guaranteed to converge with a limited generalization error and does not suffer from overfitting problem due to *the Law of Large Numbers* [31].

**Figure 3.**
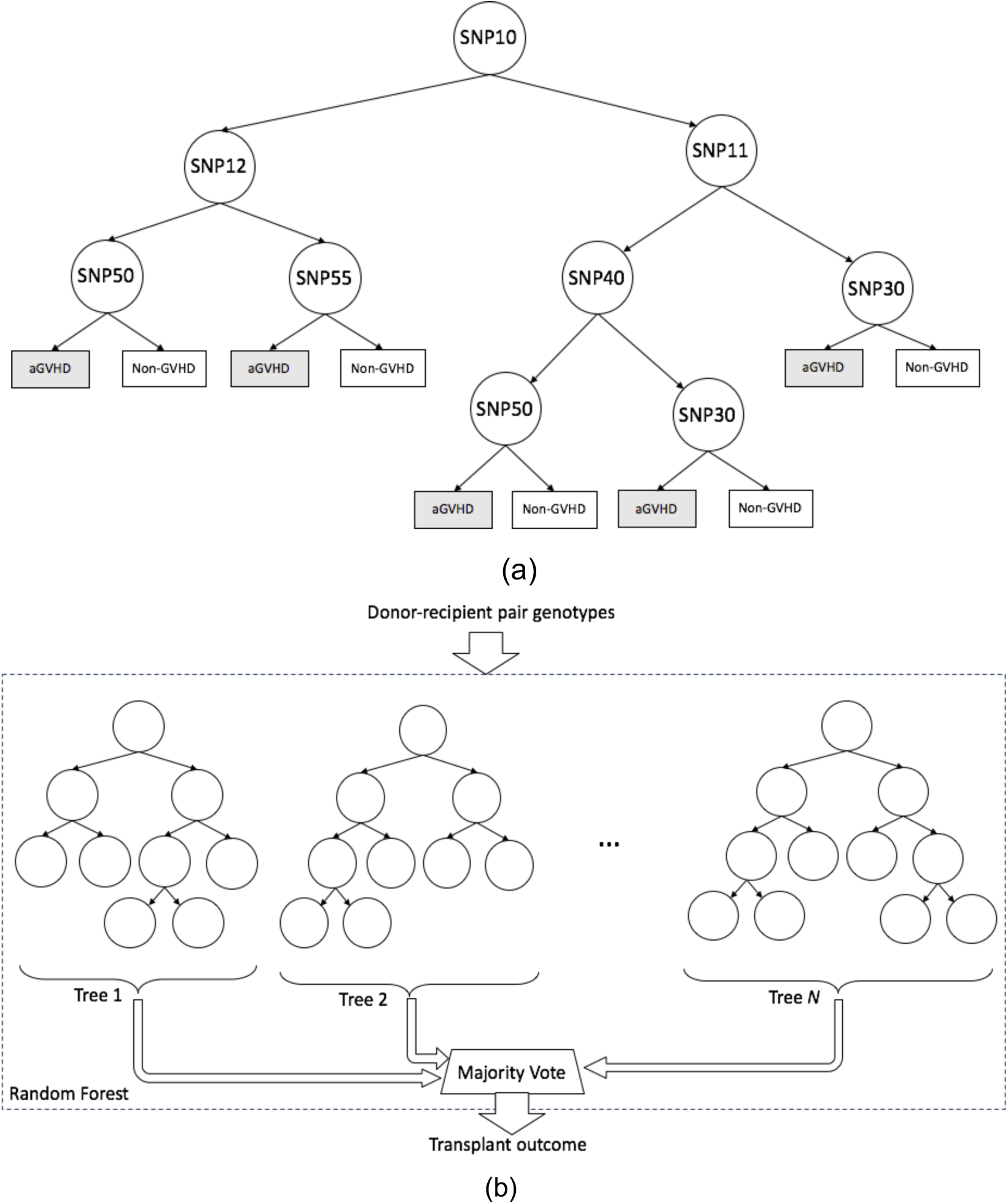
Diagram of single decision tree and the random forests. (a) a single decision tree in the forest; (b) Random Forest classifying transplant outcome from the donor-recipient pair genotype*s*.

In addition to its effective predictive ability, RFs also measure the importance of the variables in terms of their relevance to the phenotypic outcome. This function has shown great potential in genome-wide association studies and bioinformatic applications due to its effectiveness and potential interpretability. The original RF measures the feature importance using two different metrics.

The first variance importance metric is called Gini importance (GIMP). At a node in a tree, the objective is to reduce the class ambiguity as the tree grows and the split at a node is determined by the feature that reduces the class ambiguity the most when the sample passes down the split. In RF, the impurity of splits is measured by the Gini impurity index [32], defined as follows: suppose at a node, observations are trained using feature set *R*_*m*_ = {*f*_*m*1_, *f*_*m*2_, …, *f*_*md*_}. Write (*x*_*i*_, *y*_*i*_) to denote each observation, where *x*_*i*_ has *d*-dimensional features and *y*_*i*_ is the corresponding outcome label of *K* possible classes, *y*_*i*_.∈ {1,2,…,*K*} The frequency of class *k* at node *m* is defined as

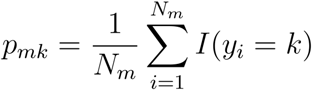

where 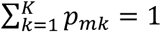. The final class of the observation at the node is determined as *p*_*mk*_, i.e., the majority class in the node *m*. For binary classification (*K* = 2), the Gini impurity index is defined as

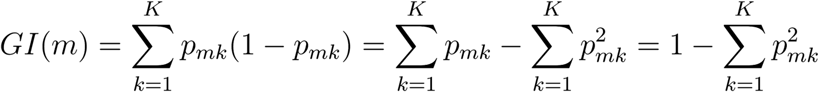

In our case-control cases, there are two classes: AML patient as 1 and healthy donor as 0 for scenario 1; or acute GVHD group as 1 and non-acute GVHD group as 0 for scenario 2. In both cases, the Gini index is

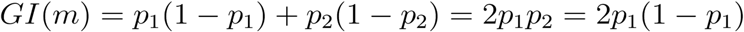

where *p*_1_ and *p*_2_ are the probabilities of the two classes mentioned above, respectively, and *p*_1_ + *p*_2_ = 1.

The Gini importance (GIMP) of a feature in a tree is calculated as the sum of the Gini impurity decrease from a parent node to its children nodes over all nodes in the tree. The GIMP score in the RF is defined as the sum (or average) of the Gini importance value among all trees in the forest.

The second feature importance is based on the feature’s predictability. After estimating the OOB prediction error during the training phase, the feature values in the OOB data set are randomly permuted and fed into the trained RF. The difference between the OOB prediction error and the permuted prediction error is defined as the prediction-based feature importance. If this value is a large positive value, the corresponding feature has high predictability and is favored high in the ranking; whereas negative or zero values indicate the features are not predictive and thus are discarded in ranking.

It has been shown that both of these metrics suffer a certain degree of selection bias when ranking features. The GIMP favors the features with many possible split points, i.e., categorical variables with many categories or continuous variable [33]. In genomic variance selection, it tends to be in favor of SNPs with high minor allele frequencies (MAF)[34, 35]. Many studies have proposed correction methods to eliminate bias. Altmann et al. [36] proposed to permute the response (phenotypic outcome) to calculate the null importance distribution while preserving the relationships between features. The algorithm is shown to reduce the feature selection bias induced by the GIMP but also provides the significance level *P*-values for each feature. Later, Janitza et al. Janitzza et al. [37] proposed an alternative approach to improve the computational speed while correcting the feature selection bias and providing the *P*-values for each feature. Nembrini et al. [38] provided a unified framework with a corrected impurity importance measure (AIR) to calculate the GIMP fast and they claimed that AIR outperforms the previous approaches in terms of computational performance and statistical power. All these bias correction methods have been incorporated and implemented in the R package `ranger` [39], and the Altmann-corrected GIMP is adopted in this study.

The prediction-based importance (PIMP) does not have these issues; however, it tends to favor the features that locate closer to the root node since they tend to affect the prediction accuracy of a larger set of observations and the permutation-based importance favors these variables [33]. A modified PIMP was proposed by Ishwaran [40], where it follows the same procedure as in the original RF, except instead of permuting the features in the out-of-bag data and test on the trained trees from the in-bag data, here the trees are randomized by using left-right random daughter assignment at each of the features. When a case is dropped down to the node with the feature in question, the left and the right daughter nodes of the following lower trees are chosen randomly with the same probability to till it reaches the leaf node. This procedure promotes the poor leaf node values for cases that pass through the nodes that split on the feature.

The predictability of the selected feature set is assessed by using OOB samples with the overall classification error, area under the receiver operating characteristic curve (AUC), and the normalized Brier score defined by Ishwarn and Lu [41]. Brier score is more stable than AUC when assessing the classifier performance. A value of 100 normalized Brier score indicates random guessing and 0 being a perfect classifier.

Figure 4 shows the proposed iRBA-RF feature selection model. During the first stage, noise and phenotypically irrelevant features are removed through the iRBA using MultiSURF as its core RBA. By removing the lowest ranked features, it retains the multi-way interaction relationships between features from MultiSURF. The refined feature set is then fed into the RF model in the second stage. The RF then train models and rank the features through GIMP and/or PIMP metrics. In this study, we implemented the model by incorporating the `scikit-rebate` library written in Python [28] (available at https://github.com/EpistasisLab/scikit-rebate) and two random forest R packages, `ranger` [39] and `randomForestSRC` [41, 42].

**Figure 4.**
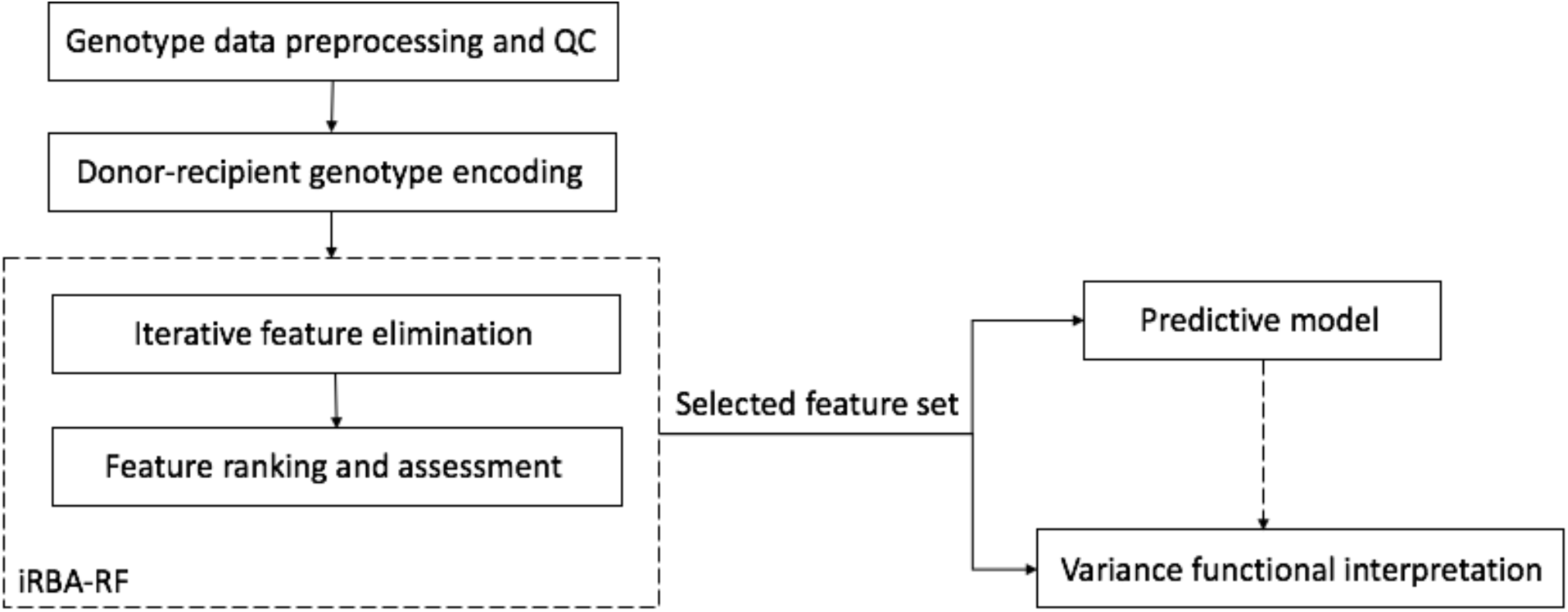
Illustration of iRBA-RF feature selection model

## Data Collection and Preprocessing

A retrospective cohort of blood cancer patients and their HLA matching donors have been selected in this study. The microarray genotype data collection and primary analysis have been described in [43]. In order to reduce the bias induced by disease types and the reference population, we chose AML patients and their transplant cases and used the original genotypes without imputation. After data quality control [supplementary material 7], 331 transplant cases (662 individuals in total) of AML patients and HLA matching donors with 630,793 genotyped autosomal SNPs were included in this study. SNPs from the sex chromosome were excluded from this study; however, sex-mismatch conditions were considered as clinical characteristics in Scenario 2 acute GVHD case-control context.

As described in the Methods section, we investigated the iRBA-RF model in two scenarios. In Scenario 1, the formatted genotype matrix has a size of 662×630,793 and the AML disease status as its target label; in Scenario 2, the formatted genotype matrix has a size of 311×261,586 and the acute GVHD status as the target label.

## Results

### Scenario 1: AML case-control experiment

The original 630,793 SNPs were reduced to 200 SNPs through the iRBA-RF and they were further reduced to 176 SNPs and 164 SNPs using GIMP and PIMP, respectively. Table 1 shows the top 30 SNPs ranked by the GIMP scores with their significance *P*-values. Of the 176 GIMP-based SNPs, 103 SNPs showed statistically significant scores at the confidence level *α* = 0.05. The full list for the 200 SNPs can be found in the Supplementary Table 1.

**Table 1.** Top 30 SNPs linked to AML, which are ranked by the Gini impurity importance using the bias-corrected metric. For illustration purpose, here lists the top 30 SNPs out 200 SNPs from the proposed feature selection model.

| Rank | Marker | CHR:POS | Gene(s) | Major | Minor | MAF | Importance score | $p$-values |
| --- | --- | --- | --- | --- | --- | --- | --- | --- |
| 1 | rs2694642 | chr2:61369045 | *USP34* | A | G | 0.315 | 1.669 | 0.010 |
| 2 | rs928770 | chr21:38265545 | *KCNJ15* | C | T | 0.287 | 0.932 | 0.010 |
| 3 | rs10936248 | chr3:161818648 |  | C | T | 0.383 | 0.854 | 0.020 |
| 4 | rs4698732 | chr4:14336718 |  | C | T | 0.442 | 0.832 | 0.010 |
| 5 | rs4692262 | chr4:27923209 | LOC105374552* | C | A | 0.372 | 0.770 | 0.040 |
| 6 | rs11869908 | chr17:72674911 | SLC39A11 | G | T | 0.210 | 0.768 | 0.010 |
| 7 | rs7718156 | chr5:172669363 | NEURL1B | C | A | 0.321 | 0.719 | 0.010 |
| 8 | rs2293836 | chr14:79840947 | NRXN3 | T | C | 0.112 | 0.694 | 0.010 |
| 9 | rs6915644 | chr6:65106919 | EYS | G | A | 0.397 | 0.692 | 0.010 |
| 10 | rs10002187 | chr4:149994262 | DCLK2** | G | A | 0.190 | 0.689 | 0.010 |
| 11 | rs749773 | chr2:165877228 | TTC21B | T | C | 0.199 | 0.644 | 0.020 |
| 12 | rs285206 | chr20:43667902 | MYBL2 | T | C | 0.202 | 0.605 | 0.010 |
| 13 | rs10926025 | chr1:239949314 | LOC105373224 | C | T | 0.298 | 0.597 | 0.010 |
| 14 | rs12675334 | chr8:83887031 |  | A | G | 0.396 | 0.587 | 0.020 |
| 15 | rs9819506 | chr3:172452314 | GHSR*;<br>BZW1P1**;<br>TNFSF10***;<br>FNDC38*** | C | T | 0.427 | 0.578 | 0.030 |
| 16 | rs6106323 | chr20:2169032 | STK35**;<br>LOC105372502** | G | A | 0.257 | 0.574 | 0.010 |
| 17 | rs2914290 | chr5:7629643 | ADCY2 | C | T | 0.181 | 0.565 | 0.010 |
| 18 | rs1365342 | chr4:37097911 | LOC101928721 | G | A | 0.329 | 0.564 | 0.040 |
| 19 | rs1173099 | chr9:90679392 | DIRAS2**;<br>OR7E109P*** | T | G | 0.260 | 0.556 | 0.010 |
| 20 | rs2222514 | chr7:123206473 | SLC13A*;<br>LYPLA1P1** | A | G | 0.432 | 0.548 | 0.010 |
| 21 | rs12714359 | chr2:2603252 | LOC105373389***;<br>LOC107985839*** | A | G | 0.367 | 0.548 | 0.020 |
| 22 | rs2185591 | chr20:43133279 | *PTPRT* | C | A | 0.423 | 0.542 | 0.020 |
| 23 | rs788871 | chr1:30591356 | *MATN1* *** | T | C | 0.474 | 0.528 | 0.030 |
| 24 | rs17329514 | chr18:71998994 | LOC102725148***;<br>LOC105372189*** | A | G | 0.101 | 0.522 | 0.010 |
| 25 | rs41135 | chr5:96830323 | *ERAP1* | G | A | 0.369 | 0.519 | 0.020 |
| 26 | rs428148 | chr2:70583509 | *TGFA* *; *ADD2* *** | C | T | 0.298 | 0.518 | 0.030 |
| 27 | rs10794031 | chr10:125876751 | *DHX32* | A | G | 0.439 | 0.517 | 0.020 |
| 28 | rs725856 | chr4:39746658 | *UBE2K* | A | G | 0.124 | 0.510 | 0.010 |
| 29 | rs675992 | chr1:17888266 | *ACTL8* *** | A | G | 0.255 | 0.492 | 0.040 |
| 30 | rs2025009 | chr14:68376888 | *RAD51B* | C | G | 0.476 | 0.491 | 0.020 |

\*: genes that are within 10 kb range of upstream or downstream from the marker
\*\*: genes that are outside 10 kb but within 50 kb range from the marker
\*\*\*: genes that are outside 50 kb but within 100 kb range from the marker

The PIMP scores are further assessed through the delete-*d* Jackknife subsampling scheme as proposed in [41]. Figure 5 illustrates the 95% asymptotic normal confidence intervals for the top 50 SNPs ranked by the median PIMP scores. For a full list of features by PIMP, please refer to Supplementary Table 2. Compared to the SNPs listed in Table 1, 9 SNPs (rs2694642 (*USP34*), rs928770 (*KCNJ15*), rs10936248, rs2293836 (*NRXN3*), rs6915644 (*EYS*), rs1173099, rs788871, rs17329514, rs675992) are ranked in the top 30 in both cases, whereas 3 SNPs (rs10002187, rs6106323, rs1365342) from Table 1 ranked between 31 and 50 in Figure 5.

**Figure 5.**
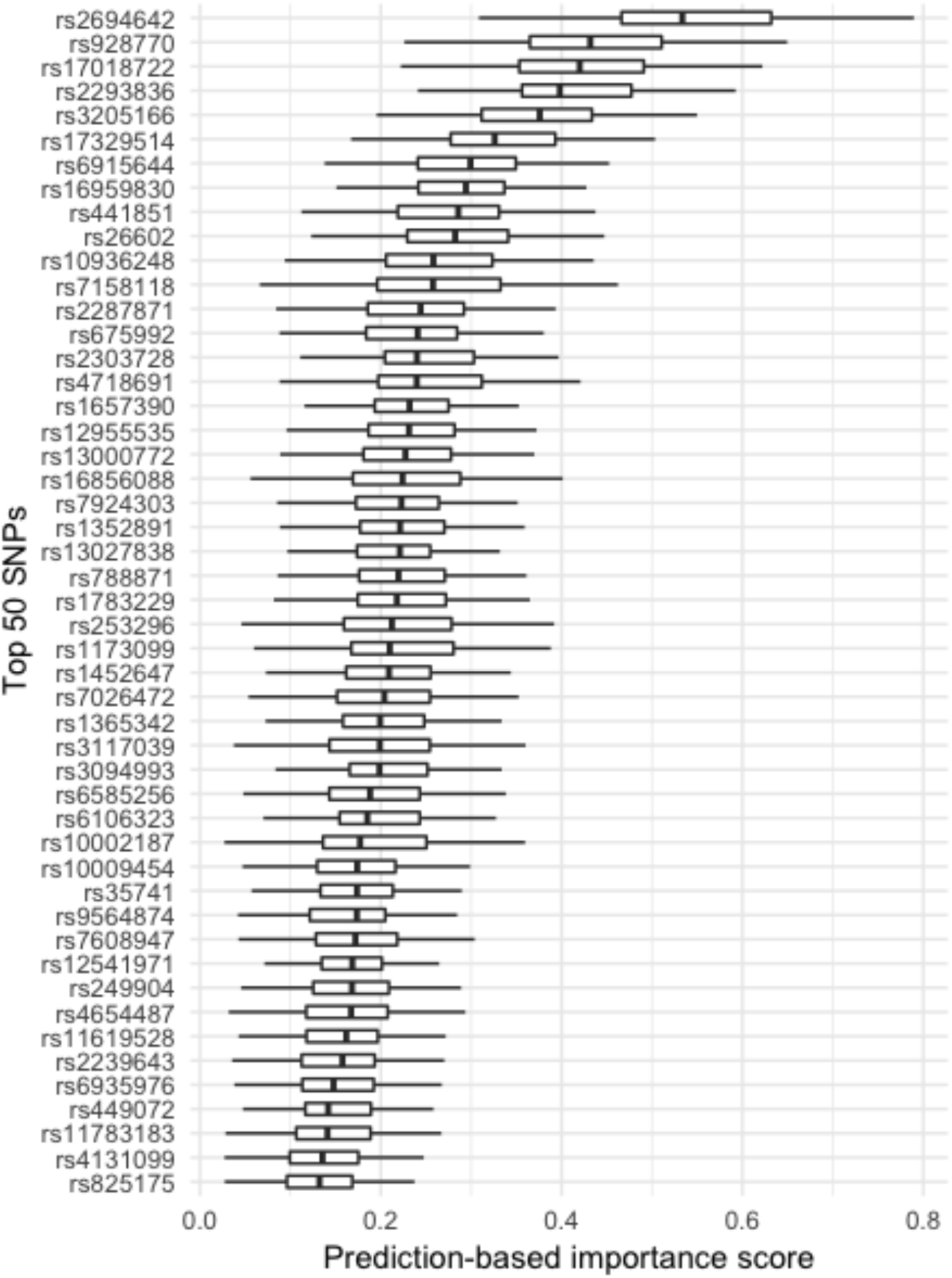
Delete-*d* Jackknife 95% asymptotic normal confidence intervals for the top 50 SNPs in the AML case-control scenario. The large positive variance importance values indicate the high predictability of the features, whereas zero and negative values suggest noise variables.

### Scenario 2: acute GVHD case-control

In the case of acute GVHD, the genotype matrix has twice as many dimensions as Scenario 1, since the donor and recipient genotypes were concatenated in the same vector for each case. The original genotype matrix has a total of 1,261,586 SNPs, and after the iRBA-RF, the number was reduced to 400 SNPs. The classical HLA typing (HLA-A, -B, -C, -DQB1, -DRB1) and DR sex mismatch status are major factors that influence the transplant outcome, and hence these two types of variables were added to the reduced genotype matrix before RF feature ranking. A total of 411 variables (400 SNPs, 10 HLA gene typing, 1 sex mismatch status) were ranked through the RF using GIMP and PIMP metrics, respectively.

342 out of 411 variables were selected by GIMP, only 124 of which showed statistically significant scores at the confidence level *α*=0.05. Top 30 variables by GIMP is listed in Table 2, and the full list can be found in Supplementary Table 3. Similar to Scenario 1, PIMP scores are assessed through delete-*d* jackknife subsampling procedure and estimated the 95% asymptotic normal confidence interval. 297 variables were selected through PIMP scores, top 50 of which are shown in Figure 6, and the full list of PIMP features can be found in Supplementary Table 4.

**Table 2.** Top 30 variables linked to acute GVHD, which are ranked by the bias-corrected Altmann-GIMP. For illustration purposes, here lists the top 30 variables out 411 SNPs from the iterative feature selection model.

| Rank | Marker | CHR: POS | Gene(s) | Major | Minor | MAF | Source | Importance score | P-value |
| --- | --- | --- | --- | --- | --- | --- | --- | --- | --- |
| 1 | rs10936748 | chr3:173283597 | *LOC105374224* | T | G | 0.151 | recipient | 0.562 | 0.020 |
| 2 | rs2389923 | chr4:119810542 | *LINC01365\*\** | A | G | 0.246 | donor | 0.507 | 0.010 |
| 3 | rs17172094 | chr7:42622588 | LOC105375251 | G | A | 0.435 | recipient | 0.474 | 0.010 |
| 4 | rs3818283 | chr9:27169126 | TEK | C | T | 0.252 | recipient | 0.473 | 0.010 |
| 5 | rs4262322 | chr8:14839455 | SGCZ | G | T | 0.244 | donor | 0.472 | 0.020 |
| 6 | rs1410267 | chr13:97614111 | LOC105370324 | A | C | 0.306 | donor | 0.471 | 0.010 |
| 7 | rs17161332 | chr7:78675562 | MAGI2 | T | C | 0.069 | recipient | 0.417 | 0.010 |
| 8 | rs1341852 | chr13:60350653 | LOC105370228 | A | G | 0.449 | recipient | 0.416 | 0.010 |
| 9 | rs7940835 | chr11:3244721 | MPGPRES** | T | C | 0.140 | recipient | 0.408 | 0.010 |
| 10 | rs7187289 | chr16:67933975 | PSMB10*; CTRL*;<br>PSKH1*;<br>LCAT*;<br>SLC12A4* | A | C | 0.320 | donor | 0.406 | 0.010 |
| 11 | rs4638670 | chr18:27701183 | LOC105372042*** | A | C | 0.174 | recipient | 0.404 | 0.010 |
| 12 | rs354843 | chr4:141267650 | ZNF330**;<br>RNF150*** | T | C | 0.269 | donor | 0.400 | 0.010 |
| 13 | rs719910 | chr7:42722647 | LINC01448** | T | C | 0.399 | recipient | 0.395 | 0.010 |
| 14 | rs6461551 | chr7:21312333 |  | G | A | 0.383 | recipient | 0.391 | 0.010 |
| 15 | rs17236893 | chr3:170745833 | LOC101928583 | A | G | 0.095 | recipient | 0.367 | 0.010 |
| 16 | rs10974006 | chr9:38738297 |  | G | T | 0.257 | recipient | 0.364 | 0.020 |
| 17 | rs273592 | chr11:30857302 | LOC107984419 | A | G | 0.482 | recipient | 0.361 | 0.010 |
| 18 | rs2481955 | chr13:28009444 | FLT3 | G | A | 0.378 | donor | 0.360 | 0.010 |
| 19 | rs227130 | chr20:8452312 | PLCB1 | G | A | 0.370 | recipient | 0.350 | 0.020 |
| 20 | rs11160228 | chr14:95051806 | DICER1** | G | A | 0.242 | recipient | 0.336 | 0.010 |
| 21 | rs4863533 | chr4:137992400 | LOC107986315**;<br>LOC105377447**;<br>LINC00616**;<br>SLC7A11*** | G | A | 0.270 | recipient | 0.333 | 0.010 |
| 22 | rs2868956 | chr19:28320511 | LOC107985269** | T | C | 0.235 | recipient | 0.329 | 0.020 |
| 23 | rs13035654 | chr2:139894716 |  | T | C | 0.182 | recipient | 0.328 | 0.020 |
| 24 | rs509012 | chr13:21720524 | FGF9** | G | A | 0.225 | donor | 0.316 | 0.020 |
| 25 | rs10503960 | chr8:34340069 | RPL10AP3*** | A | C | 0.181 | donor | 0.314 | 0.010 |
| 26 | rs12203592 | chr6:396321 | IRF4 | C | T | 0.037 | recipient | 0.304 | 0.010 |
| 27 | rs12763563 | chr10:12989971 | CCDC3 | G | A | 0.220 | recipient | 0.301 | 0.010 |
| 28 | rs4685366 | chr3:16614824 | DAZL* | A | G | 0.445 | donor | 0.301 | 0.030 |
| 29 | rs504371 | chr6:165310563 | C6orf118;<br>LOC105378113 | G | T | 0.431 | recipient | 0.298 | 0.010 |
| 30 | rs2161495 | chr5:103750446 | LOC105379107 | C | T | 0.313 | recipient | 0.297 | 0.010 |

\*: genes that are within 10 kb range of upstream or downstream from the marker
\*\*: genes that are outside 10 kb but within 50 kb range from the marker
\*\*\*: genes that are outside 50 kb but within 100 kb range from the marker

**Figure 6.**
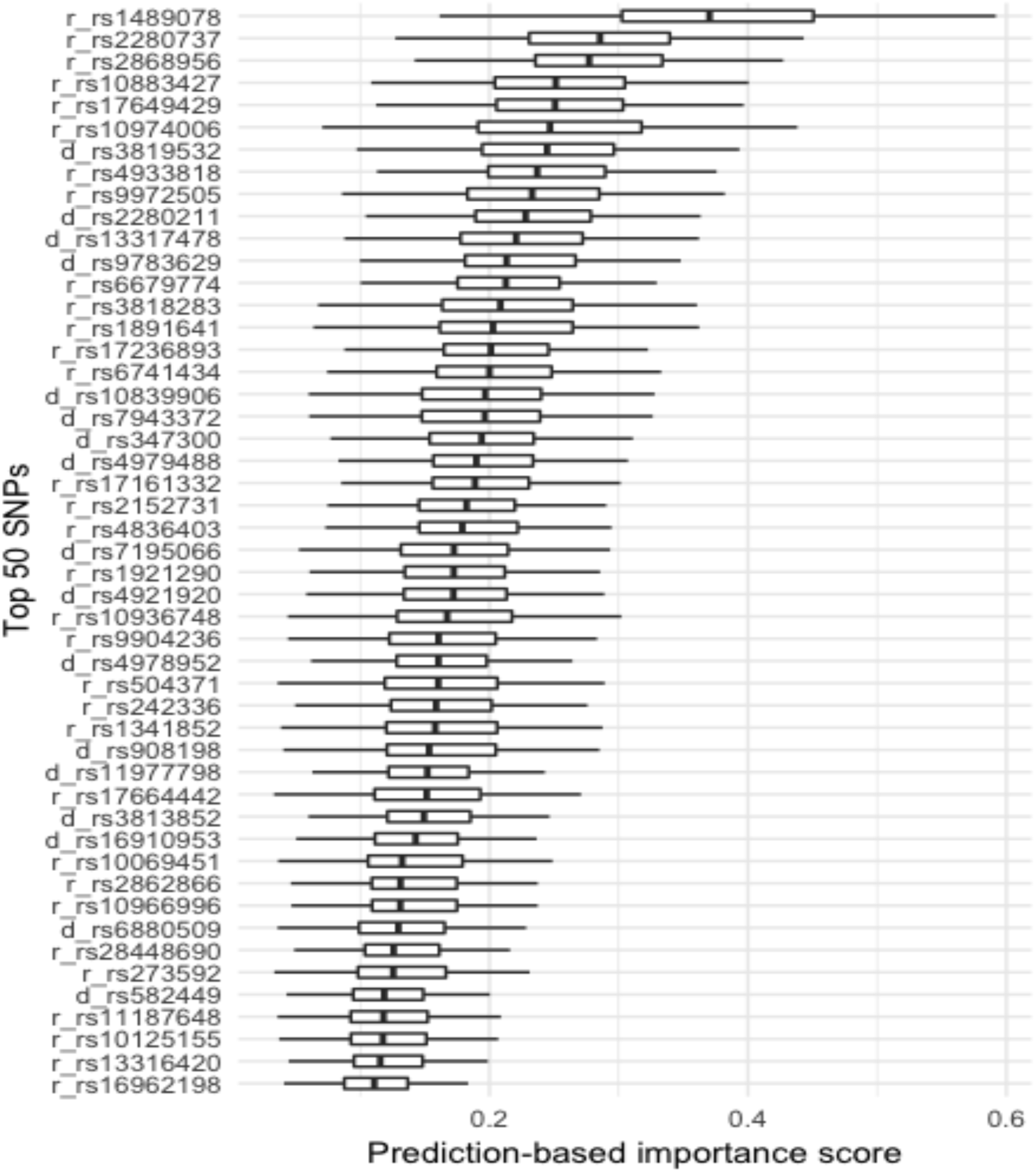
Delete-*d* Jackknife 95% asymptotic normal confidence intervals for the top 50 SNPs in the acute GHVD case-control scenario. The large positive variance importance values indicate the high predictability of the features, whereas zero and negative values suggest noise variables. ‘r_’ indicates SNPs from recipients, while ‘d_’ for SNPs from donors.

Compared to GIMP features in Table 2, 6 SNPs [rs10936748 (*LOC105374224*), rs3818283 (*TEK*), rs17161332 (*SGCZ*), rs17236893 (*LOC101928583*), rs10974006, rs2868956] are ranked in the top 30 in both cases, whereas 4 SNPs [rs1341852 (*LOC105370228*), rs11160228, rs4863533, rs504371 (*C6orf118*)] from Table 2 ranked between 31 and 50 in Figure 5.

## Discussions

Ideally, the features that are selected by iRBA-RF have the best predictability on the phenotypic outcomes and the related gene regions are actively involved in the pathways of the diseases in question. To assess the results, we first examined the predictability of the selected feature sets by comparing the classification performance to a random set of features with the same size. The classification performance was examined in three criteria: the normalized Brier score where a score of 100 indicates a random guess and the lower, the better classifier performance, the AUC, and the overall OOB error rate. Figure 7 shows the comparison among different feature sets. The random feature sets (Random200/Random400) were selected 1000 times, while the rest feature groups were trained and tested on OOB samples 1000 times. The selected features through iRBA-RF (Top200/Top400, GIMP, PIMP) in both scenarios show significantly superior classification performance in all three criteria (*p* < 2.2e-16). Within the selected groups (Top200/Top400, GIMP, PIMP), the pairwise t-tests showed a significant difference between each group using all three criteria (*p* < 0.001), except for the Brier scores between Top200 and GIMP groups. As shown in Figure 7, the classifiers using the PIMP-based feature sets generally showed better predictive performance, and this is mainly because PIMP-based features are ranked based on the classifier’s performance.

**Figure 7.**
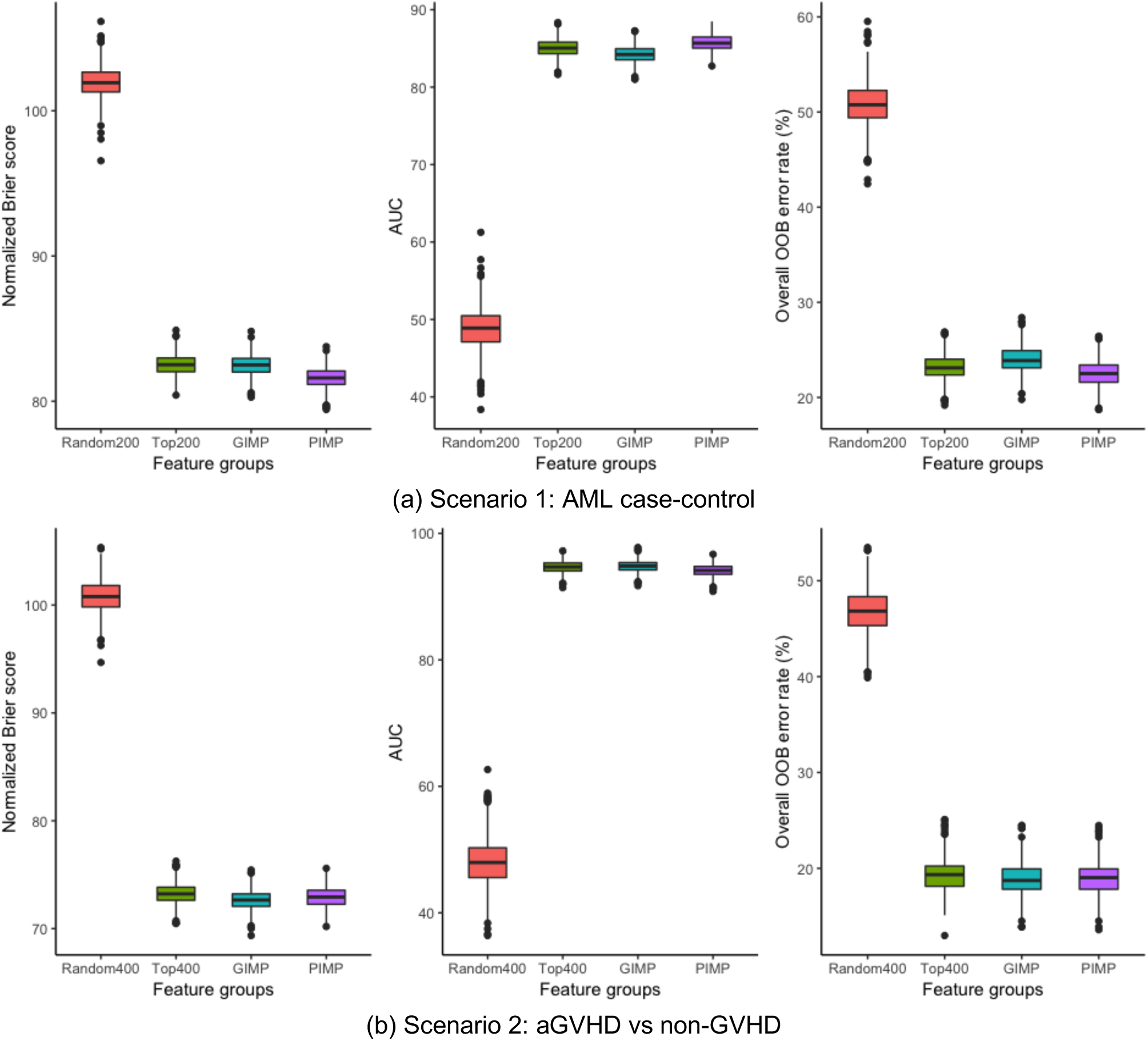
(a) Scenario 1: AML case control. (b) Scenario 2: aGVHD vs non-GVHD. Comparison of four different sets of features: (1) Random200 (or Random400): 200 (or 400) features randomly selected from the original feature set. (2) Top200 (or Top400): top ranking 200 (or 400) features selected by the iRBA-RF algorithm (3) GIMP: 176 (or 342) SNPs out of the selected 200 (or 400) SNPs using the GIMP score, (4) PIMP: 164 (or 297) features out of the selected 200 (or 400) SNPs using the PIMP score. Each feature sets were trained 1000 times and evaluated by the normalized Brier scores, AUC and overall error rate of OOB samples, respectively.

From the functional point of view, the top-ranked features are not random and evidence can be found in the literature. In Scenario 1, multiple SNPs among the selected 200 SNPs are from the following functional gene groups that are reported to be linked to AML [44, 45]. These gene groups include spliceosome (rs10794031, rs3205166), cohesin complex (rs2025009), epigenetic modifiers (rs1987193), serine/threonine kinase (rs10002187, rs994502, rs13000880), protein tyrosine phosphatases (rs2185591) and other myeloid transcription factors (rs285206). Most of these SNPs didn’t show significant importance scores, however, *SLC39A11* (rs11869908), *MYBL2* (rs285206), *PTPRT* (rs2185591), *DHX32* (rs10794031), *RAD15B* (rs2025009) are ranked the top 30 GIMP with *p*-value<0.05.

*SLC39A11* (rs11869908), also known as *ZIP11*, is a zinc transporter gene that has been reported to be linked to multiple cancers [46]. Specifically, a high expression of *ZIP4* and low expression of *ZIP11* are significantly associated with the higher grade of Glioma[47]. In addition, mutations in *IDH1* is reported to be highly correlated with higher expression of *ZIP11*, suggesting a possible synergistic interaction between *IDH1* and *ZIP11* [47]. *MYBL2* (rs285206) has an essential role in cell cycle progression, cell survival, and cell differentiation, and is found to be overexpressed in multiple cancer cases [48]. Overexpression of *MYBL2* is suggested to have a prognostic value for disease-free survival and cumulative incidence of remission for AML patients [48, 49]. *PTPRT* (rs2185591) encodes cellular signaling proteins that regulate cell growth, differentiation, and oncogenic transformation and is reported to be in the genetic interaction network of AML mutational landscape [50–52]. *DHX32* is an RNA helicase that is reported to associate with the acute lymphoblastic leukemia [53, 54]. Notably, its dysregulation is believed to contribute to carcinogenesis [55]. *RAD51B* (rs2025009) encodes one of the *RAD51* paralogs that participate in DNA repair, and the polymorphisms in the gene and the gene inactivation through chromosome translocation have been demonstrated to be linked to AML, breast cancer, head and neck cancer [56–59].

Other genes ranked high in the list have also shown evidence of roles in AML linked pathways. For instance, the top-ranked gene *USP34* (rs2694642) is reported to regulate the levels of axin and stabilize beta-catenin and further modulate Wnt signaling pathway positively [60]. Wnt/*β*-catenin pathway has shown to be essential in AML for leukemia stem cells to develop and thus allow malignant progression [61–66]. *KCNJ15* (rs928770) encodes potassium inwardly-rectifying channel on the cell membrane and is reportedly a susceptible gene for Type 2 diabetes [67, 68] and linked to the hematological traits and clinical features of Down syndrome [69–71]. *NEURL1B* is a paralog of *NEURL1*, and the deletion of this gene region has been linked to adult *de novo* AML [44].

In the case of acute GVHD, most of the top-ranked SNPs do not lie in a gene region; however, they are within 50 kb range of downstream or upstream of the coding genes. Interestingly, multiple gene regions from donors are ranked high in both GIMP- and PIMP-based feature set, suggesting the potential role of genetic polymorphisms from graft stem cells in the transplant outcomes. Genes that are linked to abnormalities of skin and the gastrointestinal (GI) tract are also selected by the iRBA-RF algorithm, all of which are the main symptoms of acute GVHD.

Notably, rs7187289 (donor) is located on chromosome 16q22.1, where five genes (*PSMB10, CTRL, PSKH1, LCAT, SLC12A4*) tightly clustered together [72]. It is in the upstream of *PSMB10*, an immunoproteasome subunit that plays a vital role in major histocompatibility complex (MHC) class I restricted antigen processing and presentation [73], T-cell polarization and differentiation, and cytokine production by macrophages [74]. Hence, *PSMB10* is believed to involve in the development of inflammatory autoimmune diseases and hematologic malignancies and to be a marker of cell damage and immunological activity [75]. In renal transplantation, it has recently reported being associated with chronic antibody-mediated rejection (AMR) and posed as a potential intragraft and peripheral blood marker of acute rejection [76, 77]. The impairment of immunoproteasome subunits is critical for malignant cells to escape immune recognition, suggesting its possible role in the graft-versus-tumor effect after allo-HCT. LCAT is secreted by the liver and generally believed to maintain the unesterified cholesterol gradient between peripheral cells and high-density lipoprotein (HDL)[78]. *SLC12A4* encodes a human potassium chloride cotransporter 1 (KCC1)[79], and the dysfunction of the membrane ion channels has been reported to link to several diseases, like sickle cell disease [80].

Several gene regions in the list have direct roles in the signs of acute GVHD in the GI tract. *CTRL* is a chymotrypsin-like protease expressed in the pancreas and secreted in pancreatic juice [81] and is well known to be downregulated in pancreatic cancer [82]. PSKH1 protein is mainly found in Golgi apparatus, endoplasmic reticulum (ER), nucleus, cell membrane, cytoskeleton [83] and believed to play a role in intranuclear serine/arginine-rich domain (SR protein) trafficking and pre-mRNA processing [84]. A recent study suggested it possibly linked to the pathogenesis of Crohn’s disease [85]. *MAGI2* (rs17161332) encodes a scaffolding protein that involved in epithelial integrity, and studies have shown that the genetic variation in *MAGI2* is linked to the inflammatory bowel disease (IBD), i.e., ulcerative colitis and Crohn’s disease [86]. Moreover, *IRF4* (rs12203592) controls T_H2_ (T helper cell) responses and intestinal Th17 cell differentiation, mucosal cytokine IL-17 regulation, suggesting a central of *IRF4* in immune regulation in the gut [87–92].

The risk of getting a secondary solid cancer following allo-HCT is substantially higher than in general population, and the risk factors have been well documented [93–95]. Melanoma, breast cancer, thyroid cancer, prostate cancer, and cervix cancer are the most frequently occurring cancer types after allo-HCT in the recipient’s later life. The genes that were selected in this study have evidence to link to the cancer progression and may be able to explain the incidents. The feature sets after iRBA-RF include multiple genes that play critical roles in the progression of carcinogenesis. For example, *SGCZ* (rs4262322) encodes a protein that plays a role in maintaining cell membrane stability and have been reported its role linked to cancer development and progression [96]. *DICER1* (rs11160228) encodes essential proteins for a micro-RNA processing pathway and plays a central role in epigenetic modulation of gene expression, and downregulation of DICER1 expression has been reported to be linked to a wide range of cancer types [97, 98].

A few leukocyte-specific genes, such as *TEK, FLT3*, and *PLCB1* are also ranked high in the list. *TEK* (rs3818283) encodes angiopoietin-1 receptor that is critical to the induction and growth of new blood vessels and influence tumor growth. It has been reported that mutations in *TEK* are linked to AML suggesting its essential role in leukemogenesis depending on an uncharacterized cellular context [99, 100]. A more recent study demonstrated that angiogenesis precedes leukocyte infiltration during inflammation suggesting the essential involvement of angiogenesis in the initiation of inflammatory diseases, such as acute GVHD and IBD [101]. Proteins encoded by *FLT3* (rs2481955) stimulates hematopoiesis and is reportedly expressed at high levels in a spectrum of hematologic malignancies, including AML. Mutations in FLT3 usually lead to a poor prognosis and is suggested to be a potential therapeutic target for kinase inhibitor [102]. Similarly, *PLCB1* (rs227130) is recently proposed as a potential therapeutic target for AML patients, since the monoallelic deletion or increased PLCB1 expression is a prognostic factor and is reportedly linked to the transition from myelodysplastic syndromes (MDS) to AML [103–105].

These genes and their functional interpretation seem to explain potential underlying mechanisms; however, they by no means explain the actual biological functions nor do they provide a full picture of the genetic interactions in AML or acute GVHD. The SNPs used in this study are common polymorphisms (MAF>0.005), and the locations are much sparser than the whole genome sequences. Thus, the representation of genes from the SNPs is merely a remote approximation. The highly ranked SNPs are not necessarily directly involved in the pathways linked to the disease; instead, some ungenotyped genes that share a high linkage disequilibrium with those SNPs may exert more significant influence on the disease status. On the other hand, the feature interactions captured through iRBA-RF suggest the statistical epistasis among these features or SNPs but not the biological epistasis. Therefore, functional interpretation from SNP sets needs much careful consideration. More rigorous experiments may be needed to validate the potential genetic interactions. Overall, it is a promising start to investigate the genetic interactions in transplant-related outcome studies while considering both donor’s and recipient’s genomes simultaneously.

One of the advantages of using GIMP and PIMP-ranked features in RF is that it removes the arbitrariness of choosing the number of features. Positive values of GIMP or PIMP indicate the features contribute positively to the predictive power of the predictive models and a value of zero is an appropriate cutoff. The *p*-values of GIMP scores add additional confidence level to the selection, as do the confidence intervals to the PIMP scores. On the other hand, GIMP ranking and PIMP ranking are not always consistent with each other, as they are using two different criteria to measure the feature importance. Moreover, as many researchers have point out [106, 107], a *p*-value is not a reliable metric to infer the significance, since it depends on assumptions of the models and the sample size and lacks reproducibility. Lu et al. [108] proposed standard error and confidence intervals of PIMP as an alternative to the *p*-values of regression models and demonstrated the robustness of PIMP to the sample size and model assumptions. Especially, when the sample size is small, it is a more reliable indicator than *p*-values. Therefore, it is desirable to use the PIMP ranked feature set for further downstream analysis.

There are several parameters to determine for the iRBA-RF model. There are three parameters for the first step of feature elimination using iRBA: the optimal subset size (*Ns*), the number of iterations (*Iteration*), the percentage of features that will be removed after each iteration (*pct*), the number of features to keep after the last iteration (*featNum*). The original VLSReliefF algorithm suggested a large sample size and relatively small *Ns* achieve reliable results[30]. By default, top 50th percentile (*pct*=0.5) rank of SNPs is selected after each iteration. However, in our experiment, this removes many interactive and relevant variables, and the final feature sets have very little predictive power. The goal of iRBA is to remove as many irrelevant variants as possible while keeping all possible interacting ones. In this study, we chose the following parameters for both scenarios: *Ns*=1000, *Iteration*=5, *pct*=0.25. As for *featNum*, we employed a grid search strategy to find the optimum values for each of the scenarios. As shown in Figure 8, *featNum*=200 for Scenario 1 and *featNum*=400 for Scenario 2 achieved the best classification performance, measured by the normalized Brier score, AUC and overall OOB error rate. As for RF models, each forest has 300 trees (*ntree*=300) with the default *mtry*=sqrt(dimension).

**Figure 8.**
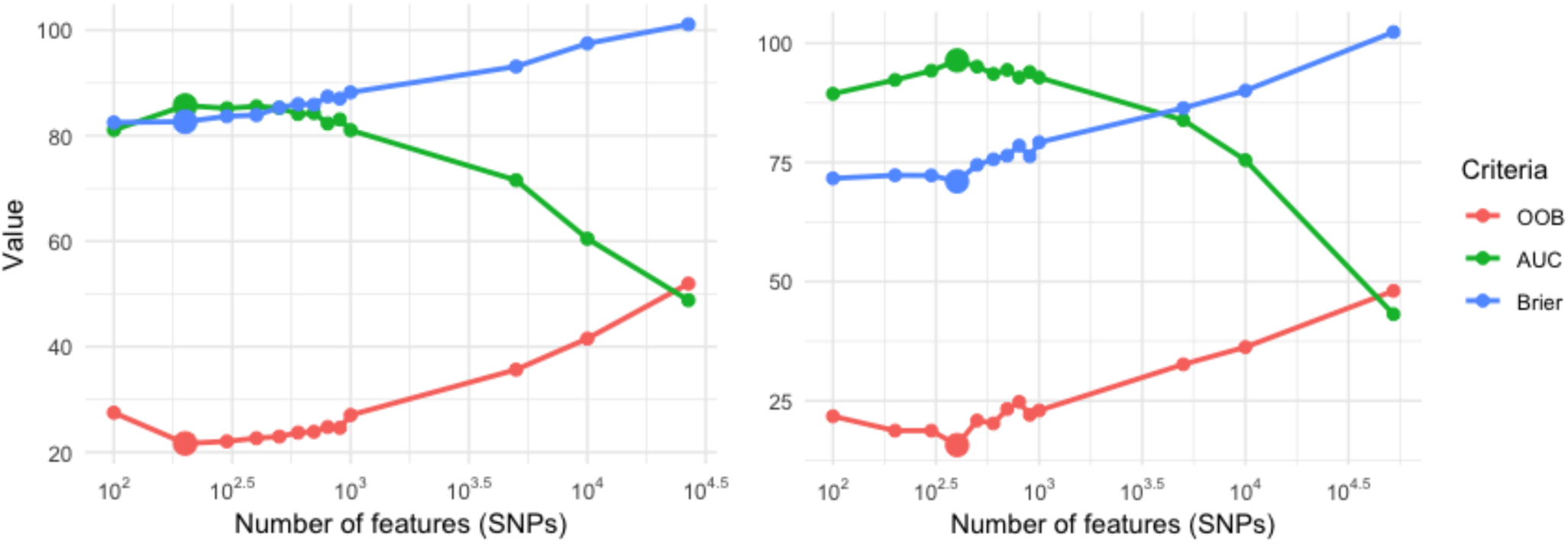
The effect of the number of SNPs. from 10,000, 5000, 1000, 900, 700, 500, 300, 200, 100: the OOB prediction error reaches its minimum between 300 and 500 SNPs for both AML and acute GVHD. Number of features are shown on a log_10_ scale. OOB: Out-of-bag sample average prediction error, shown in percentage (%); AUC: Area under the ROC; Brier: the normalized Brier score. The optimal feature size would produce the minimum OOB error rate (%), the minimum Brier score and the maximum AUC value.

One caveat of the iRBA-RF model, however, is that, unlike regression or a simple associations test, it cannot tell the direction of a variable’s impact on the disease (positive or negative, protective or progressive) from the model. On the other hand, this may avoid or minimize the effect of Simpson’s paradox, where the positive or negative association of variables reverse sign due to the change of a confounding factor. Simpson’s paradox is a common issue in association studies, especially in a high-dimensional bioinformatics data set [109]. Keep in mind that, the selected features collectively contribute to the predictive power and thus it is not an indication of the influence of a single SNPs on the disease. It is worth noting that the importance scores and feature ranking are a relative concept, and they have little indication of biological importance. In summary, the iRBA-RF model offers a computationally efficient and functionally effective method to find the candidate variable groups whose interactions may exert to the disease status. With a larger cohort size with broader genomic coverage, it may effectively find the genetic interaction networks that are directly linked to the disease development.

## Conclusion

We developed a hybrid feature selection model, iRBA-RF, to reduce the feature space and ultimately rank and select the variants that may be linked to the diseases in question, AML, and acute GVHD. The proposed model successfully selected the most related SNPs out of over 600 K and 1 M SNPs and produced a reasonable predictive accuracy. The model was applied to genomic data in this study, but it can be extended to examine multi-omics data with other clinical characteristics, as well as the multi-class prediction problems.

As discussed above, evidence of the genes can be found in the literature to be linked to the disease in question; however, in order to determine their biological role and further assist optimized donor selection process and personalized therapeutic development, experiments on a larger cohort size, along with immunological wet lab validation experiments on the selected genes, are desired.

## Acknowledgement

The authors thank Dr. Abeer Madbouly from the Center for International Blood and Marrow Transplant Research (CIBMTR) for curating and providing the data used in this study.

## Supplementary Tables 1-4

[*Supplementary tables.xlsx*]

Supplementary Table 1. Top ranking SNPs from GIMP for Scenario 1: AML case-control

Supplementary Table 2. Top ranking SNPs from PIMP for Scenario 1: AML case-control

Supplementary Table 3. Top ranking SNPs from GIMP for Scenario 2: acute GVHD case-control

Supplementary Table 4. Top ranking SNPs from PIMP for Scenario 2: acute GVHD case-control

**Supplementary Table 5.**
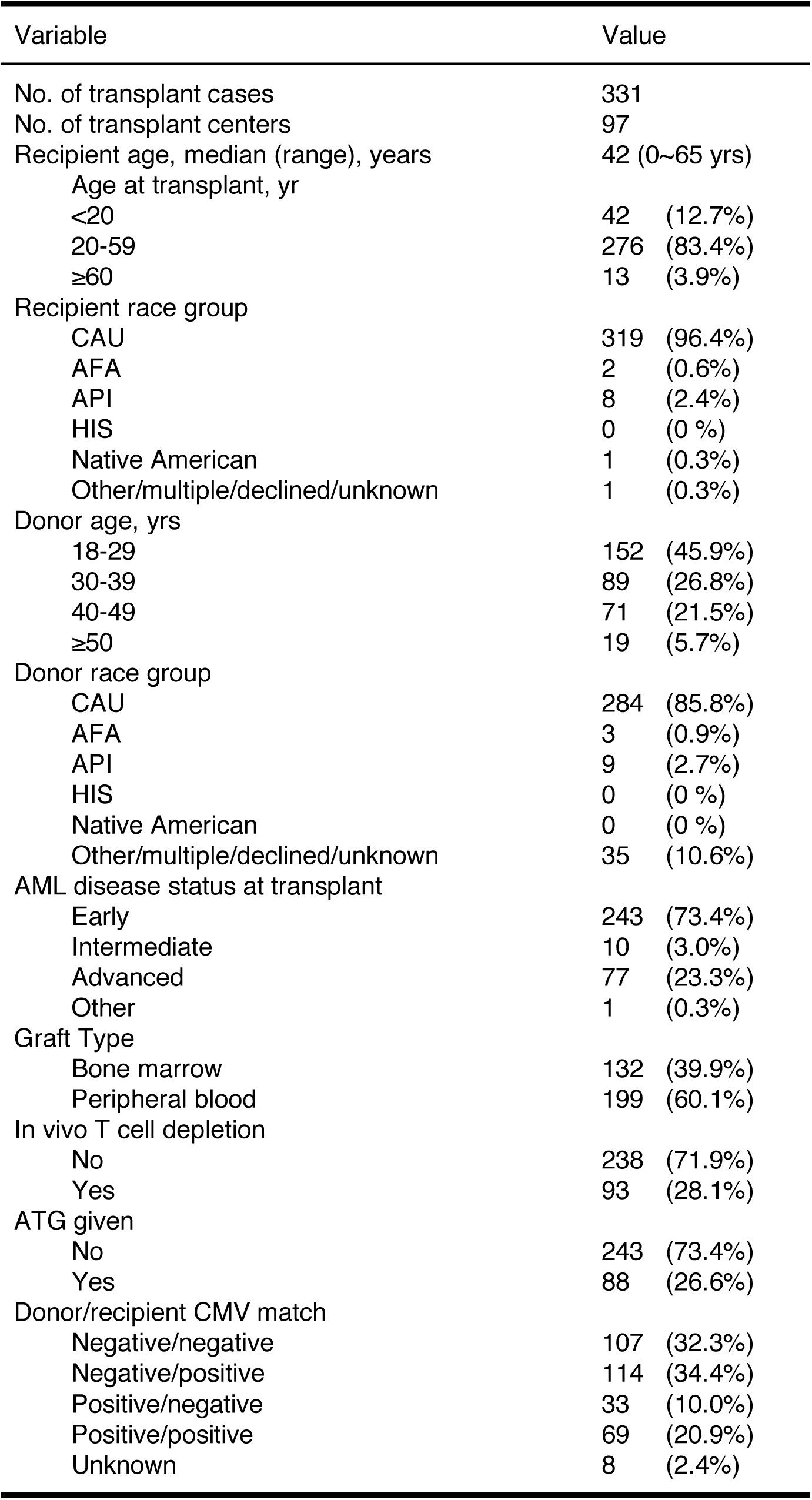
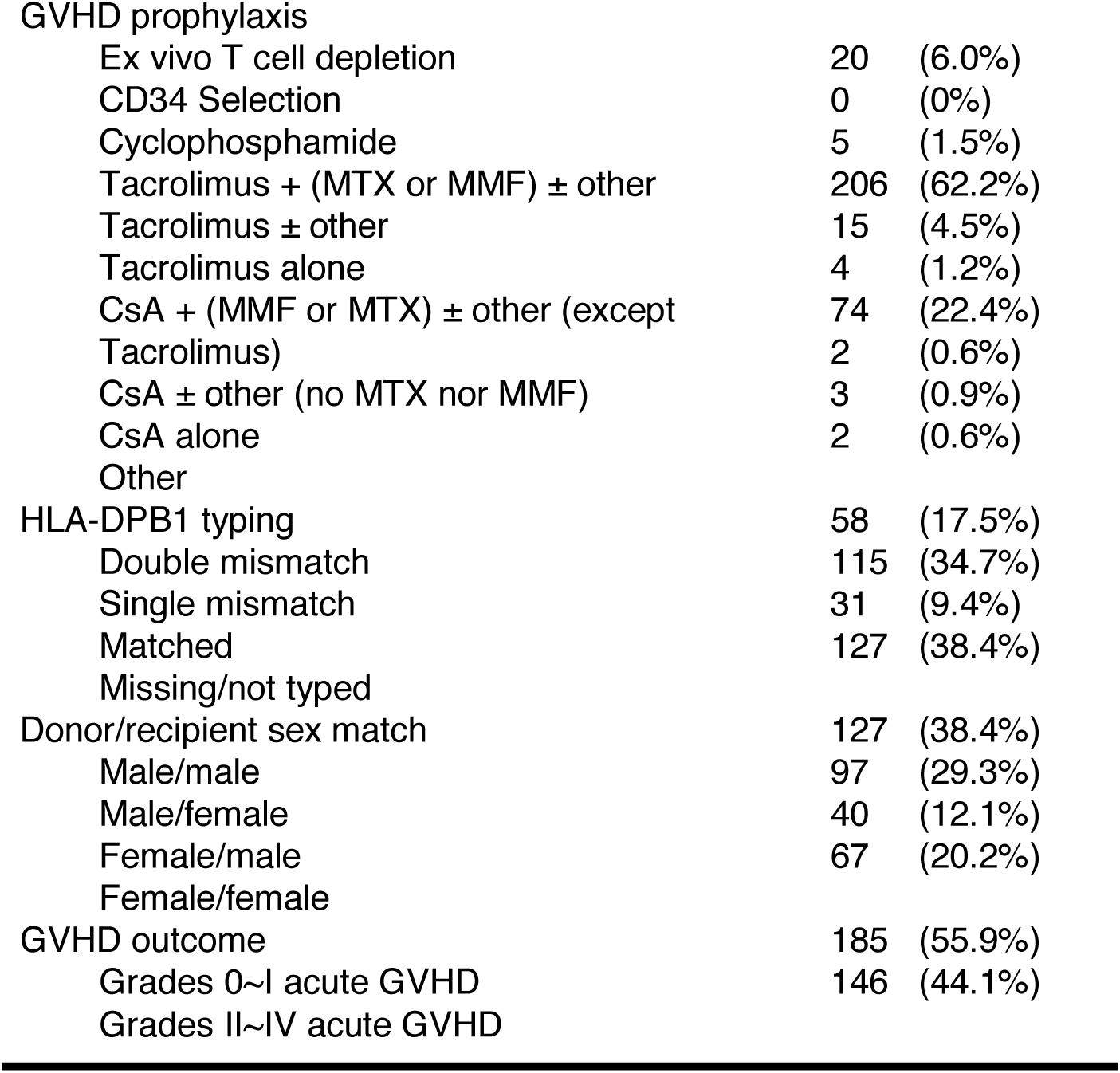
Characteristics of acute myeloid leukemia patients transplant cases

**Supplementary table 6.** HLA typing summaries

| HLA-A | Freq | HLA-B | Freq | HLA-C | Freq |
| --- | --- | --- | --- | --- | --- |
| A*02:01 | 174 | B*07:02 | 114 | C*07:01 | 126 |
| A*01:01 | 115 | B*08:01 | 107 | C*07:02 | 115 |
| A*03:01 | 94 | B*44:02 | 73 | C*05:01 | 71 |
| A*24:02 | 50 | B*40:01 | 41 | C*03:04 | 56 |
| A*11:01 | 32 | B*15:01 | 40 | C*06:02 | 56 |
| A*29:02 | 21 | B*35:01 | 37 | C*04:01 | 54 |
| A*68:01 | 21 | B*18:01 | 36 | C*03:03 | 29 |
| A*26:01 | 18 | B*27:05 | 21 | C*12:03 | 26 |
| A*32:01 | 17 | B*13:02 | 19 | C*08:02 | 20 |
| A*31:01 | 15 | B*44:03 | 19 | C*01:02 | 19 |
| A*25:01 | 13 | B*57:01 | 16 | C*02:02 | 16 |
| A*23:01 | 12 | B*14:02 | 15 | C*16:01 | 12 |
| A*30:01 | 12 | B*38:01 | 13 | C*07:04 | 10 |
| A*33:01 | 6 | B*37:01 | 11 | C*12:02 | 8 |
| A*68:02 | 6 | B*52:01 | 8 | C*14:02 | 3 |
| A*02:05 | 3 | B*49:01 | 7 | C*17:01 | 2 |
| A*30:02 | 3 | B*51:01 | 6 | C*02:10 | 1 |
| A*03:02 | 2 | B*55:01 | 6 | C*08:01 | 1 |
| A*24:03 | 1 | B*56:01 | 6 | C*15:02 | 1 |
| A*24:17 | 1 | B*14:01 | 5 |  |  |
| A*30:04 | 1 | B*40:02 | 5 |  |  |
| A*36:01 | 1 | B*35:03 | 3 |  |  |
|  |  | B*39:01 | 3 |  |  |
|  |  | B*41:01 | 3 |  |  |
|  |  | B*45:01 | 3 |  |  |
|  |  | B*50:01 | 3 |  |  |
|  |  | B*15:17 | 2 |  |  |
|  |  | B*27:02 | 2 |  |  |
|  |  | B*35:02 | 2 |  |  |
|  |  | B*47:01 | 2 |  |  |
|  |  | B*53:01 | 2 |  |  |
|  |  | B*58:01 | 2 |  |  |
|  |  | B*15:02 | 1 |  |  |
|  |  | B*15:03 | 1 |  |  |
|  |  | B*15:10 | 1 |  |  |
|  |  | B*15:18 | 1 |  |  |
|  |  | B*35:08 | 1 |  |  |
|  |  | B*58:02 | 1 |  |  |

| HLA-DRB1 | Freq |
| --- | --- |
| DRB1*15:01 | 116 |
| DRB1*03:01 | 105 |
| DRB1*07:01 | 74 |
| DRB1*01:01 | 60 |
| DRB1*04:01 | 56 |
| DRB1*13:01 | 37 |
| DRB1*11:01 | 30 |
| DRB1*13:02 | 24 |
| DRB1*04:04 | 23 |
| DRB1*08:01 | 13 |
| DRB1*11:04 | 12 |
| DRB1*12:01 | 12 |
| DRB1*01:02 | 11 |
| DRB1*14:01 | 11 |
| DRB1*01:03 | 10 |
| DRB1*15:02 | 8 |
| DRB1*04:02 | 4 |
| DRB1*09:01 | 4 |
| DRB1*13:03 | 4 |
| DRB1*04:07 | 3 |
| DRB1*10:01 | 3 |
| DRB1*16:01 | 3 |
| DRB1*04:03 | 2 |
| DRB1*04:05 | 2 |
| DRB1*11:03 | 2 |
| DRB1*08:02 | 1 |
| DRB1*08:04 | 1 |
| DRB1*11:02 | 1 |
| DRB1*12:02 | 1 |
| DRB1*13:04 | 1 |
| DRB1*13:05 | 1 |
| DRB1*14:02 | 1 |
| DRB1*16:02 | 1 |

| HLA-DQB1 | Freq |
| --- | --- |
| DQB1*06:02 | 116 |
| DQB1*02:01 | 106 |
| DQB1*03:01 | 103 |
| DQB1*05:01 | 77 |
| DQB1*02:02 | 59 |
| DQB1*03:02 | 48 |
| DQB1*06:03 | 37 |
| DQB1*06:04 | 21 |
| DQB1*03:03 | 18 |
| DQB1*04:02 | 15 |
| DQB1*05:03 | 11 |
| DQB1*06:01 | 8 |
| DQB1*05:02 | 4 |
| DQB1*06:09 | 3 |

## Supplemental Material 7: Microarray genotype data preprocessing

The original SNP genotype data was obtained and processed by Madbouly et al. and described in (Madbouly et al. 2017). In this study, we performed the quality control separately and did not use the imputed genotypes.

We have removed individuals that show ambiguous sex genotype than their reported sex. The rest of parameters used in the SNP filtering is as follows. 1) If a SNP minor allele frequency (MAF) is less than 0.005 or showed up in less than 10 individuals, then those SNPs are filtered out. 2) SNPs that have less than 95% call rate are removed. 3) It is recommended to use the control-only samples for Hardy-Weinberg equilibrium (HWE) test (Anderson et al. 2010). Here we used the healthy donor-only samples and excluded the SNPs that have P-values lower than 0.001 after the HWE test. After these steps, we obtained 630,793 SNPs for the 331 donor-recipient pairs.

